# Acute stress impairs reward learning in men

**DOI:** 10.1101/2020.07.13.200568

**Authors:** Joana Carvalheiro, Vasco A. Conceição, Ana Mesquita, Ana Seara-Cardoso

## Abstract

Acute stress is ubiquitous in everyday life, but the extent to which acute stress affects how people learn from the outcomes of their choices is still poorly understood. Here, we investigate how acute stress impacts reward and punishment learning in men using a reinforcement-learning task. Sixty-two male participants performed the task whilst under stress and control conditions. We observed that acute stress impaired participants’ choice performance towards monetary gains, but not losses. To unravel the mechanism(s) underlying such impairment, we fitted a reinforcement-learning model to participants’ trial-by-trial choices. Computational modeling indicated that under acute stress participants learned more slowly from positive prediction errors — when the outcomes were better than expected — consistent with stress-induced dopamine disruptions. Such mechanistic understanding of how acute stress impairs reward learning is particularly important given the pervasiveness of stress in our daily life and the impact that stress can have on our wellbeing and mental health.

## 1. Introduction

Learning to choose options that lead to rewards and to avoid those that result in punishments is crucial for adaptive behavior. Situational factors, such as stress, can have deleterious effects on the ability to make the best choices and learn from them (Porcelli & Delgado, 2017). Stress is present in our day-to-day life, but, notably, how acute stress affects reward and punishment learning remains largely unknown. A growing body of evidence suggests that acute stress impairs reward-seeking behavior (Berghorst, Bogdan, Frank, & Pizzagalli, 2013; Bogdan, Perlis, Fagerness, & Pizzagalli, 2010; Bogdan, Santesso, Fagerness, Perlis, & Pizzagalli, 2011; Bogdan & Pizzagalli, 2006; Ehlers & Todd, 2017; Morris & Rottenberg, 2015; Paret & Bublatzky, 2020; but see Lighthall, Gorlick, Schoeke, Frank, & Mather, 2013), but less is known about the impact of acute stress on punishment-avoidance behavior (Aylward et al., 2019; Petzold, Plessow, Goschke, & Kirschbaum, 2010). More critically, there is even less evidence on the mechanisms that underlie the behavioral effects of acute stress on reward and punishment learning. Here, we use a computational reinforcement-learning framework to investigate the impact of acute stress on reward and punishment learning in men.

In the past decades, the use of computational modeling approaches to describe behavior-brain relationships in healthy humans has played an influential role on cognitive science (Daw & Frank, 2009; Frank, 2015; Huys, Maia, & Frank, 2016; Maia & Frank, 2011; Maia, 2015). Computational models, such as reinforcement-learning models, are built and implemented to capture very specific cognitive and neural mechanisms, thus linking different levels of analysis, from cognitive and behavioral phenomena to neurobiological mechanisms (Chater, 2009; Daw & Frank, 2009; Frank, 2015; Nair, Rutledge, & Mason, 2020). Reinforcement-learning models are considered extremely useful tools to investigate the neural computations underpinning cognition and behavior (Collins & Frank, 2013; Daw, 2011; Daw & Frank, 2009; Huys et al., 2016; Maia & Frank, 2011; Nair et al., 2020), and can thus provide a mechanistic framework to disentangle the effects of acute stress on reward and punishment learning (Aylward et al., 2019; Huys, Pizzagalli, Bogdan, & Dayan, 2013; Luksys & Sandi, 2011; Otto, Raio, Chiang, Phelps, & Daw, 2013; Radenbach et al., 2015; Robinson, Overstreet, Charney, Vytal, & Grillon, 2013).

According to reinforcement-learning theory, individuals learn to gradually select more and more often the actions that maximize rewards and those that minimize punishments by learning the values of the executed actions, and such learning is driven by prediction errors (Maia & Frank, 2011; Schultz, Dayan, & Montague, 1997; Sutton & Barto, 1998). Specifically, prediction errors — which signal the difference between obtained and expected outcomes — are used to progressively update the values of the executed actions (Collins & Frank, 2013; Maia & Frank, 2011; Schultz et al., 1997; Sutton & Barto, 1998). Prediction errors can be positive or negative. Positive prediction errors occur when outcomes are better than expected leading to reward learning (Daw & Tobler, 2014; Schultz et al., 1997). Negative prediction errors occur when outcomes are worse than expected leading to punishment learning (Daw & Tobler, 2014; Schultz et al., 1997). Importantly, reinforcement-learning models can capture how quickly rewarding and punishing outcomes are integrated over time through distinct learning rates for positive and negative prediction errors, respectively (Frank, Moustafa, Haughey, Curran, & Hutchison, 2007). Conversely, blunted signaling of positive and negative prediction errors can be captured by reinforcement-learning models as reduced positive and negative learning rates, respectively.

Dopaminergic functioning plays a key role in prediction-error-based learning (Frank, Seeberger, & O’Reilly, 2004; Glimcher, 2011; Maia & Frank, 2011; Pessiglione, Seymour, Flandin, Dolan, & Frith, 2006). Prediction-error signals are known to be encapsulated in the phasic activity of dopamine neurons (Daw & Tobler, 2014; Maia & Frank, 2011; Schultz et al., 1997). Specifically, phasic bursts of dopaminergic neurons are thought to adaptively encode positive prediction errors, whereas dopamine dips have been associated with the adaptive encoding of negative prediction errors (Daw & Tobler, 2014; Maia & Frank, 2011; Schultz et al., 1997). However, phasic-dopamine responses do not seem to be always adaptive, and there is evidence that dopamine can be phasically released in an aberrant spontaneous manner (Belujon, Grace, & Grace, 2015; Maia & Frank, 2017; Sulzer, Cragg, & Rice, 2016). Studies with non-human male animals suggest that aberrant spontaneous phasic-dopamine release increases with acute stress (Anstrom, Miczek, & Budygin, 2009; Anstrom & Woodward, 2005; Valenti, Lodge, & Grace, 2011). Crucially, exaggerated, aberrant spontaneous dopamine release seems to reduce adaptive striatal phasic bursts that signal positive prediction errors (Bilder, Volavka, Lachman, & Grace, 2004; Daberkow et al., 2013; Grace, 2016; Maia & Frank, 2017; Werlen et al., 2020). Additionally, though more speculative (Maia & Frank, 2017), aberrant spontaneous dopamine release might also block the effects of dopamine dips (Frank & O’Reilly, 2006) needed to signal negative prediction errors. Still, the extent to which stress-induced aberrant spontaneous dopamine release blunts signaling of positive and/or negative prediction errors remains poorly understood.

The aim of the present work was twofold. First, to investigate how acute stress impacted behavioral performance during reward and punishment learning. Second, to inspect the computational mechanisms behind the effects of acute stress on reward and punishment learning. Given the putative roles of phasic-dopamine responses on prediction errors signaling and of acute stress on aberrant spontaneous phasic-dopamine release, we hypothesized that acute stress would impair reward learning and that such impairment would be underpinned by a decreased learning rate for positive prediction errors. More tentatively, we hypothesized that acute stress could also impair punishment learning by decreasing the learning rate for negative prediction errors.

To test those hypotheses, we used a well-established reinforcement-learning task involving monetary gains and losses (Pessiglione et al., 2006) combined with a novel acute stress manipulation. The stress manipulation consisted of exposing participants to an uncontrollable sound (i.e., participants could not put an end to it) whilst they performed the reinforcement-learning task. We chose to use a repetitive alarm sound as stressor, as alarms can promote an alertness state accompanied by acute physiological responses (Hall et al., 2016). Furthermore, uncontrollable auditory stimuli can elevate stress responses (Arguelles, Ibeas, Ottone, & Chekherdemian, 1962; Breier et al., 1987; Rylander, 2004; Westman & Walters, 1981) and may disrupt dopaminergic mechanisms (Arnsten et al., 1998) and cognitive functioning (Glass, Reim, & Singer, 1971). Whereas in studies using other well-validated stress manipulations, such as the cold pressor task (e.g., Byrne, Cornwall, & Worthy, 2019; Lighthall et al., 2013) or the Trier social test (e.g., Kruse, Tapia León, Stalder, Stark, & Klucken, 2018; Petzold et al., 2010), stress induction precedes task-solving (hence, learning), in our study we ensured that the stress induction occurred throughout the task, i.e. concurrently with learning. To check the success of the acute-stress manipulation, we collected self-report stress levels at the end of each block of the task and measured skin conductance response (SCR) rate throughout the task. Then, we inspected how acute stress altered reward and punishment learning during the reinforcement-learning task using both classical statistical analyses and computational-model-based analyses of participants’ behavioral data. For the latter analyses, we fitted participants’ choices with a previously established, biologically inspired, reinforcement-learning model (Frank et al., 2007), which has been extensively used to investigate the cognitive and behavioral impact of pharmacological manipulations and genetic variations in the dopaminergic system in humans (Diederen et al., 2017; Doll, Hutchison, & Frank, 2011; Frank & Fossella, 2011; Frank et al., 2007; Grogan et al., 2017; Rutledge et al., 2009). The fitted reinforcement-learning model allowed us to examine mechanistically the effects of acute stress on learning rates for positive and negative prediction errors. In this study, we included only male participants due to females’ hormonal-dependent variations on stress responsivity, as well as on reward and punishment learning (Diekhof, Korf, Ott, Schädlich, & Holtfrerich, 2020; Dreher et al., 2007; Ossewaarde et al., 2010).

## 2. Materials and Methods

### 2.1. Participants

Sixty-two healthy male participants (age range = 18 – 35; *M* = 21.9, *SD* = 3.7) were recruited at University of Minho, Portugal. This sample size more than doubled the size from that of a previous study where a similar probabilistic reinforcement-learning task was applied in a within-subject design (Petzold et al., 2010). We stopped data collection after achieving this sample size, and thus independently of the statistical significance of the data. Four participants were excluded from skin-conductance analyses due to poor signal quality. No participants were *a priori* excluded from any other data analyses, although we conducted confirmatory analyses excluding potential outliers to ensure that our results were not driven by extreme values.

All participants provided their informed consent before the experimental session. All experimental procedures were approved by the Ethics Committee of University of Minho.

### 2.2. Reinforcement-learning task

After a short practice (12 trials), to familiarize participants with the task timings and response keys, participants completed four blocks of an adapted version of a well-established reinforcement-learning task (Pessiglione et al., 2006) (Fig. 1). During each block of 120 trials, participants were presented with three new pairs of abstract stimuli (40 trials per pair). Each pair of stimuli was associated with a valence: one pair of stimuli was associated with gains (“gain” 0.5€/”nothing”), a second pair associated with losses (“loss” 0.5€/”nothing”), and a third pair associated with neutral, or non-financial outcomes (“look” 0.5€ /’’nothing; for a depiction of neutral trials see Fig. S1a in the Supplementary Material). The outcome probabilities were reciprocally 0.8 and 0.2 for the stimuli in each of the three pairs. On each trial, one pair was randomly presented on the screen, with one stimulus from the pair above and the other below a central fixation cross (the stimuli position was counterbalanced across trials). Participants were instructed to choose between the two visual stimuli displayed on the computer screen to maximize payoffs. Missing choices occurred when participants did not press the response keys within 2000 ms (0.38% missing choices: 40 in the stress condition and 72 in the control condition, in a total of 29760 trials across all participants) and were signaled with a “Missed” message (no other outcome was provided). Missing choices were excluded from data analyses. Participants were informed that they would be paid the amount obtained during a randomly selected block, but they all left with the same fixed amount (15€). The experiment was programmed and presented with Cogent 2000 (http://www.vislab.ucl.ac.uk/cogent.php) implemented in MATLAB R2015a (MathWorks).

**Fig. 1.**
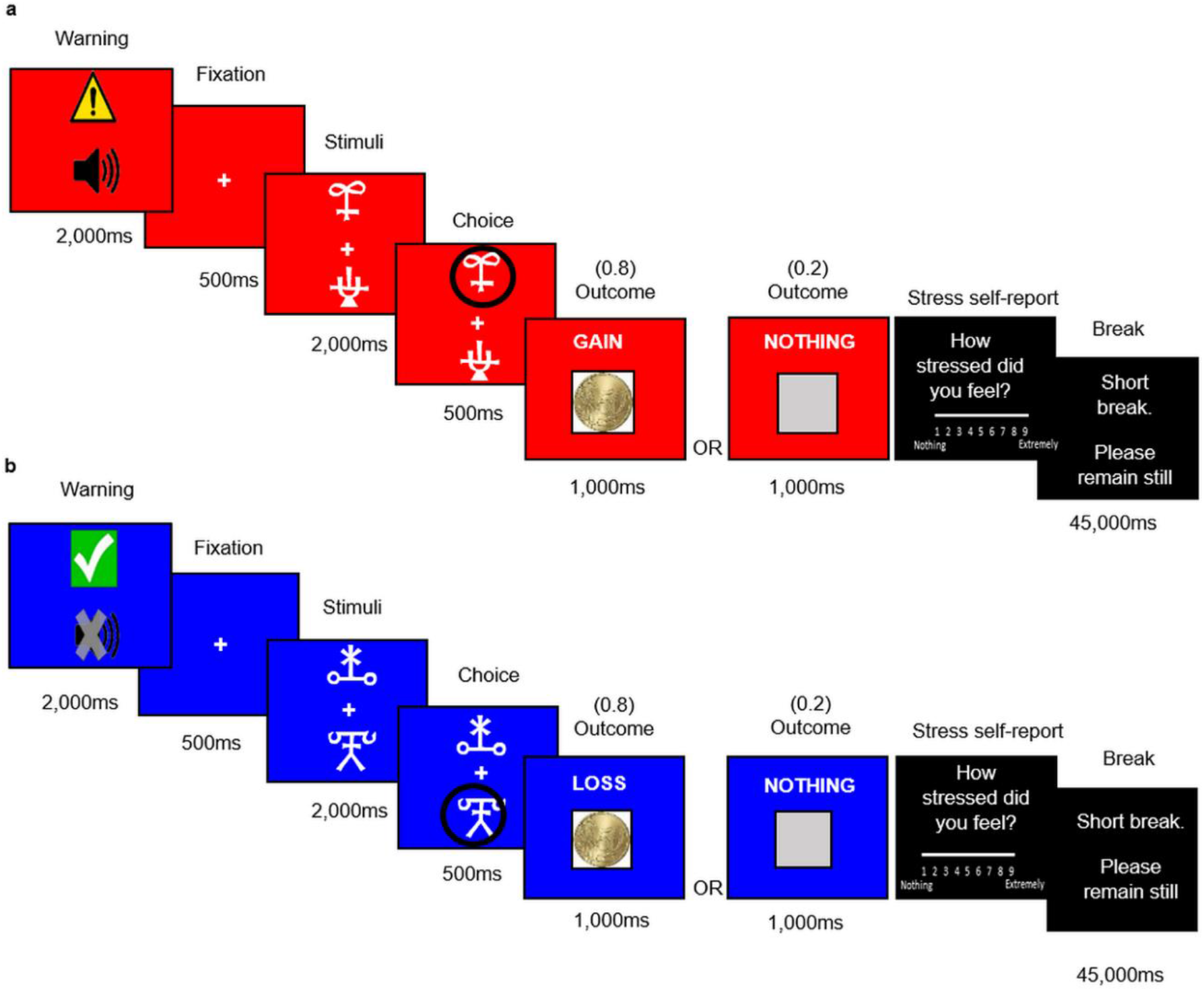
Reinforcement-learning task. On each trial of the task, participants had to choose either the upper or the lower of two abstract visual stimuli presented on a computer screen and to subsequently observe the obtained outcome, whilst under acute stress and under control conditions. **(a)** In the depicted stress-condition example, the chosen stimulus was associated with a probability of 0.8 of winning 0.5€ and with a probability of 0.2 of obtaining nothing. The other (not chosen) stimulus was associated with a probability of 0.8 of obtaining nothing and a 0.2 probability of winning 0.5€. **(b)** In the depicted control-condition example, the chosen stimulus was associated with a probability of 0.8 of losing 0.5€ and with a probability of 0.2 of obtaining nothing. The other (not chosen) stimulus was associated with a probability of 0.8 of obtaining nothing and with a 0.2 probability of losing 0.5€. Participants completed a total of four blocks, consisting of an alternation between two stress and two control blocks. To assess stress responses, self-reported stress levels were collected at the end of each block and skin conductance was measured throughout the task.

### 2.3. Acute stress manipulation

During the experimental session, participants performed two blocks of the reinforcement-learning task whilst exposed to a stressor (i.e., stress condition; Fig. 1a) and two blocks without the stressor (i.e., control condition; Fig. 1b). To elicit stress responses we exposed participants to a predictable, but uncontrollable auditory stimulus, a constant alarm (“Annoying modern office building alarm.wav”, retrieved from freesound.org, and programmed to loop uninterruptedly), played through the same set of over-ear headphones (GOODIS model GWH4093, with the volume set to the maximum), with the sound volume adjusted in the laptop to level 27 (in a scale from 0-100) for all participants. We conducted a brief pilot to qualitatively confirm that this volume was tolerable, yet stressful. Stress blocks were signaled by a warning sign and a red background (Fig. 1a), and control blocks were signaled by a safe sign and blue background (Fig. 1b). Stress and control blocks were administered alternately and in counterbalanced order. The experiment was conducted in a soundproof room to avoid interference due to environmental noise, and between 12 pm and 6 pm to minimize diurnal variability in stress responses.

### 2.4. Manipulation check

Stress levels were assessed by asking participants at the end of each block to rate how stressed they felt during that block on a scale of 1 (nothing) to 9 (extremely). To further assess the impact of the acute stressor on autonomic responses, we acquired skin conductance responses (SCRs) using BIOPAC MP150 and a pair of finger electrodes. Electrodes were attached to participants’ left index and ring fingers; the gain was set to 5, the low pass filter to 10 Hz, and the high pass filters to DC. Recordings were performed using Acqknowledge 4.4. Data were acquired at 200 Hz, downsampled to 62 Hz, and smoothed with a median filter in order to remove outliers. Each participant’s SCRs were detected using a threshold of 0 μs and a rejection rate of 10% (Kim, Bang, & Kim, 2004). The SCR rate was calculated by dividing the number of SCRs detected in each block by the duration of that block (in minutes).

Self-reported stress levels and SCR rates were analysed using repeated-measures analyses of variance (ANOVAs), with condition (stress and control) and block (1 and 2) as within-subject factors, and post-hoc paired *t*-tests. ANOVAs effect sizes are reported as eta-squared, *η^2^*, and post-hoc paired *t*-tests’ effect sizes are reported as Cohen’s *d* and 95% confidence intervals. We further conducted non-parametric Wilcoxon signed-rank tests, which are more robust to outliers, to confirm the results from post-hoc paired *t*-tests. Additionally, to confirm that our findings were robust to extreme values, we repeated the SCR rate analyses excluding participants with abnormally large SCR rates. Statistical analyses were conducted using JASP 0.9.

### 2.5. Task performance analyses

To examine the impact of acute stress on choice performance during the reinforcement-learning task, we applied a generalized linear mixed-effects (glme) model to participants’ choice data (with correct and incorrect choices coded as 1 and 0, respectively). We used a “logit” link function to account for the binomial distribution of the data. As predictor variables in the glme model we included condition (stress or control), valence (gains or losses), block number (1 or 2), and trial number (1 to 40), and the interaction of interest (condition × valence; see Table S1 in the Supplementary Material for a full description of the glme model). The glme included a fixed intercept, as well as random intercepts for each participant. We fitted the glme model to the behavioral data using MATLAB’s *fitglme* function and conducted post-hoc analyses via contrast matrices using the MATLAB’s *coefTest* function. To assess the robustness of our findings, we also tested the significance of the interaction of interest (condition × valence) using confirmatory likelihood ratio tests (Daw, 2011) between the aforementioned full glme model and two nested models, which assumed equal performance in stress and control conditions during either gain or loss trials, through the MATLAB’s *lratiotest* function. Additionally, we repeated the analyses excluding participants that performed below chance levels, which is indicative that participants did not learn to perform the task correctly and may reflect additionally non-compliance with the experimental setting.

To confirm that the choice probabilities estimated by the glme model showed a close correspondence with the actual observed choices, we used MATLAB’s *predict* function. Then, we assessed the Pearson’s correlation between the percentage of actual “correct” choices (i.e., choice of the stimuli associated with a probability of 0.8 of winning or a probability of 0.2 of losing) for each participant in each condition (averaged across blocks) and the percentage of “correct” choices as estimated by the glme model. We also performed confirmatory Spearman’s correlations, which are more robust to outliers.

### 2.6. Computational modeling

#### 2.6.1. Reinforcement-learning model

We modeled participants’ trial-by-trial behavior in the stress and control conditions using a reinforcement-learning framework (Sutton & Barto, 1998) that has been extensively used to investigate the behavioral and neural impact of pharmacological manipulations and genetic variations in the dopaminergic system in humans (Diederen et al., 2017; Doll et al., 2011; Frank & Fossella, 2011; Frank et al., 2007; Grogan et al., 2017; Rutledge et al., 2009). Importantly, the fitted model included separate learning rates for positive (*α^+^*) and negative (*α^−^*) prediction errors, to account both for the differential firing of dopaminergic neurons for positive and negative prediction errors (Daw & Tobler, 2014; Maia & Frank, 2011; Maia & Conceição, 2017) and the differential effects of dopamine onto the plasticity of the corticostriatal synapses implicated in action-value learning (Frank & O’Reilly, 2006; Maia & Frank, 2017; Maia & Conceição, 2017; Möller & Bogacz, 2019). This model also included the inverse temperature parameter, *β*, which controls the stochasticity of choice selection, or the exploration/exploitation trade-off (Daw, 2011; Sutton & Barto, 1998), as detailed below.

In the context of our experimental study, the specific reinforcement-learning model used (Frank et al., 2007) assumes that each participant gradually learns the value of choosing a given stimulus (say A or B) from a given pair of stimuli (here, “gain” or “loss” stimuli pairs, as the “neutral” pair of stimuli always yielded null monetary outcomes) as a function of the outcome that was obtained on that trial following stimulus selection. Specifically, each expected pair-stimulus value, or *Q*-value, was initialized to zero, and for each trial, *t*, within that pair of stimuli, the value of the chosen stimulus (say A was chosen) was updated according to:

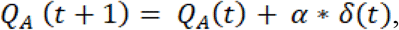

where *δ* was the prediction error:

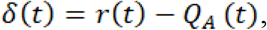

where *r*(*t*) was 0.5 for gains, 0 for neutral outcomes, and −0.5 for losses. The learning rate, *α*, was given by:

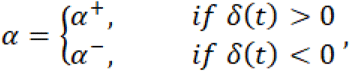

where *α^+^* and *α^−^* were the learning rates for positive and negative prediction errors, respectively (Frank et al., 2007).

The probability of choosing one stimulus over another (say A over B) was given by the *softmax* equation:

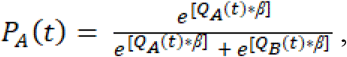

where the *β* parameter, or inverse temperature, controlled the amount of exploration/ exploitation. A null *β* results in random behavior (i.e., similar choice probabilities irrespective of the corresponding action values), whereas the higher the *β* the more deterministic is the behavior (in our case, the stimulus with the highest expected value becomes progressively more and more likely to be the one selected from the pair).

#### 2.6.2. Model fitting, parameter analyses, and model validation

We fitted the reinforcement-learning model to the trial-by-trial choice data from each participant in each condition. Model fitting involved estimating the values of the parameters (*α^+^, α^−^* and *β*) that best accounted for the respective trial-by-trial choices in each condition.

We estimated the best-fitting model parameters (*α^+^, α^−^* and *β*) for each subject in each condition using maximum *a posteriori* estimation (Daw, 2011). Specifically, to optimize model parameters, we drew the learning rates from Beta distributions [*Beta*(1.1, 1.1)] and the inverse temperature from a Gamma distribution [*Gamma*(1.2, 5)] (Palminteri, Khamassi, Joffily, & Coricelli, 2015). We then used the MATLAB’s *fmincon* function, initialized at 100 random starting points of the parameter space, to search for the parameter values that minimized the negative log posterior of the observed sequence of choices, given the previously observed outcomes, with respect to different settings of the model parameters (Daw, 2011).

To assess the effects of acute stress on the parameters estimated by the reinforcement-learning model, we conducted repeated-measures ANOVAs with condition (stress and control) and valence (positive and negative) as within-subject factors, and post-hoc paired *t*-tests. ANOVAs’ effect sizes are reported as eta-squared, η^2^, and post-hoc paired *t*-tests effect sizes are reported as Cohen’s *d* and 95% confidence intervals. We also confirmed all results from post-hoc tests using non-parametric tests (Wilcoxon signed-rank tests), which are more robust to outliers. Additionally, we repeated all analyses excluding participants that performed below chance levels to test whether the significance of the results remained unchanged. These statistical analyses were conducted using JASP 0.9.

To validate the used reinforcement-learning model, we computed trial-by-trial choice probabilities for all participants using the best-fitting set of parameters in each condition. The actual observed choices and outcomes were used to update the choice probabilities. To assess whether the choice probabilities estimated by the reinforcement-learning model (for each subject, the choice probabilities were averaged across gains or loss trials in each condition) followed the same pattern as the actual observed choices, we conducted Pearson’s correlations and confirmatory Spearman’s correlations for both conditions, using the respective mean percentages.

To further validate the robustness of our model-fitting procedure, we examined the capacity of recovering subject-condition-specific parameters using simulated datasets. Specifically, we simulated the task-choice behavior of 62 virtual participants using the parameter values that we had estimated for each of the 62 participants in each condition. We ran 100 simulations. Then, for each simulation, we fitted the model to the virtual participants’ data to estimate new (recovered) parameters. Finally, we tested the correlations between the original and these recovered parameters using Pearson’s and confirmatory Spearman’s correlations.

## 3. Results

### 3.1. Manipulation check

First, we confirmed that the acute stress manipulation successfully elicited stress responses in the participants. Self-reported stress levels differed significantly between conditions, *F*(1, 61) = 107.67, *p* < .001, *η^2^* = 0.64, as participants reported higher levels of stress in the stress condition (*M* = 5.16, *SEM* = 0.21) than in the control condition (*M* = 3.31, *SEM* = 0.20), *t*(61) = 10.38, *p* < .001, *d* = 1.32, 95% confidence interval (CI) = [1.49, 2.20] (Fig. 2a). SCR rate also differed significantly between conditions, *F*(1,57) = 20.61, *p* < .001, *η^2^* = 0.27, being higher in the stress condition (*M* = 2.89, *SEM* = 0.27) than in the control condition (*M* = 2.46, *SEM* = 0.22), *t*(57) = 4.54, *p* < .001, *d* = 0.57, 95% CI = [0.24, 0.62] (Fig. 2b). The condition × block interactions were non-significant for the self-reported stress levels, *F*(1, 61) = 0.004, *p* = .95, *η^2^* = 0, and for the SCR rate, *F*(1,57) = 0.46, *p* = .50, *η^2^* = 0.0080, suggesting that both the self-reported stress levels and the SCR rate remained stable across blocks within conditions. Furthermore, confirmatory non-parametric Wilcoxon signed-rank tests replicated the effects of acute stress on self-reported stress levels, when comparing the stress condition with the control condition, *Z* = 6.29, *p* < .001, and SCR rate, *Z* = 4.08, *p* < .001. Additionally, to check whether our results were robust to extreme values, we identified two participants with abnormally large SCR rate and reanalyzed the data without those participants. The significance of the results remained unchanged (main effect of condition: *F*(1, 55) = 19.47, *p* < .001, *η^2^* = 0.26), with higher SCR rate in the stress condition than in the control condition, *t*(55) = 4.41, *p* < .001, *d* = 0.59, 95% CI = [0.21, 0.56], even after excluding the two participants.

In sum, these results suggest that the acute stress manipulation successfully elicited stress responses.

**Fig. 2.**
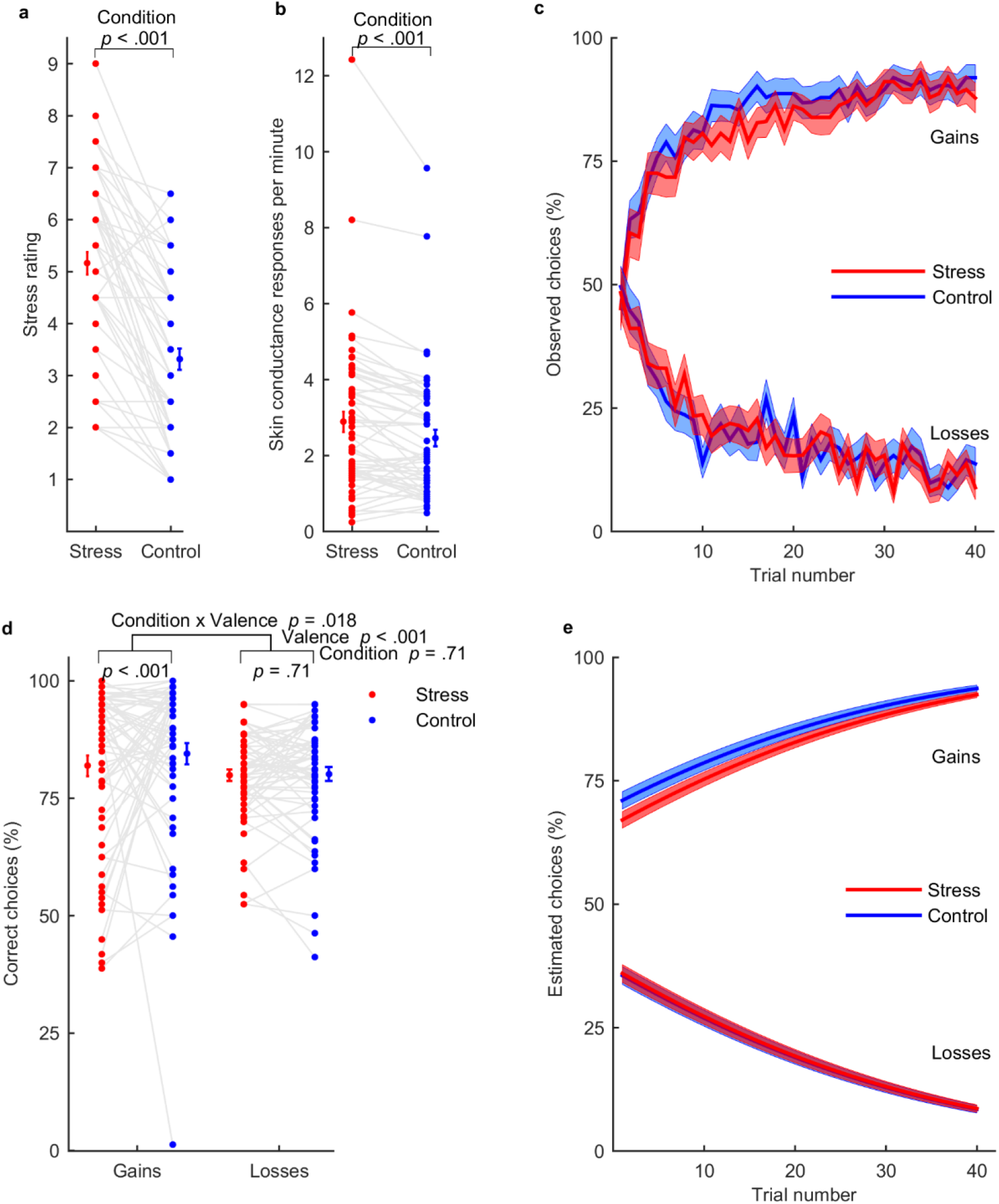
Manipulation check and task performance. **(a)** Participants (n = 62) reported higher stress levels in the stress condition (red) compared with the control condition (blue). **(b)** Skin conductance response rate (n = 58) was significantly higher in the stress condition than in the control condition. **(c)** Learning curves represent the trial-by-trial percentage of participants (n = 62) who chose the “correct” gain stimulus (associated with a probability of 0.8 of winning 0.5€; upper part of the graph) and the “incorrect” loss stimulus (associated with a probability of 0.8 of losing 0.5€; lower part of the graph), in the stress and control conditions. **(d)** Participants performed significantly worse when seeking monetary gains, but not when avoiding monetary losses, in the stress condition relatively to the control condition. The reported *p*-values are from the generalized linear mixed-effects model that included each participant’s trial-by-trial choices and respective post-hoc tests. **(e)** The choices estimated by the generalized linear mixed-effects model captured the evolution of the actual observed choices during the reinforcement-learning task (to compare the choices estimated by the model with the actual observed choices, compare the overlap of the curves between the stress and control conditions depicted here with the overlap of the curves depicted in Fig. 2c). In panels a, b, and d, connected dots represent data points from the same participant; error bars displayed on the sides of those scatter plots indicate the sample mean ± standard errors of the mean. In panels c and e, the central lines represent the means and the filled areas the ± standard errors of the means.

### 3.2. Task performance

After confirming that self-reported stress levels and SCR rate were augmented in the stress condition, we examined the impact of acute stress on choice performance during the reinforcement-learning task (Fig. 2c) using a glme model. We found a significant condition × valence interaction, *β* = −0.19, *p* = .018, 95% CI = [−0.34, −0.031] (Fig. 2d), and post-hoc analyses revealed that under stress, comparatively to the control condition, participants performed significantly worse when seeking monetary gains, *F* (1, 19755) = 12.87, *p* < .001, but not when avoiding losses, *F* (1, 19755) = 0.14, *p* = .71. As performance below chance levels might be indicative of non-compliance with the experimental setting, we also inspected whether each participant’s behavioral performance across gains and losses was below chance levels (i.e., less than 50% of correct choices) in the stress or control conditions. One participant performed below chance levels in the stress condition, and two participants performed below chance levels in the control condition. Thus, we repeated the aforementioned analyses excluding these three participants. We found that the condition × valence interaction was even more significant after excluding participants that did not learn how to perform the task, *β* = −0.28, *p* < .001, 95% CI = [−0.44, −0.12]. We also confirmed the robustness of our findings using likelihood ratio tests. We found that the full glme model (which assumed different performance towards gains between the stress and control conditions) had a significantly better fit than a model that assumed no differences in performance towards gains between conditions, χ^2^(1) = 12.80, *p* < .001, and that the same full model did not have a significantly better fit than a model that assumed no differences in performance towards losses between the stress and control conditions χ^2^(1) = 0.20, *p* = .66. These analyses indicate that behavioral performance towards gains, but not losses, significantly differed between the stress and control conditions.

In sum, acute stress selectively impaired choice performance towards monetary gains during the reinforcement-learning task. As an additional check, we confirmed that the choices estimated by the glme model showed close correspondence with the observed choices across trials in both conditions, Pearson’s *r* > 0.65, *p* < .001, Spearman’s *r* > 0.51, *p* < .001 (Fig. 2e).

Additionally, given that acute stress is thought to increase aberrant spontaneous phasic-dopamine release (Anstrom et al., 2009; Anstrom & Woodward, 2005; Valenti et al., 2011), which, in turn, may lead to augmented learning and behavioral responding for neutral stimuli (Maia & Frank, 2017; Roiser, Howes, Chaddock, Joyce, & McGuire, 2013), we performed an exploratory analysis of participants’ choices for the neutral stimuli pairs (the pairs not associated with financial outcomes; for further details see Table S2 and “Analyses of neutral trials” in the “Supplementary Analyses” section of the Supplementary Material). We found that, within the pairs of neutral stimuli, acute stress increased behavioral responding towards the stimuli that more often yielded as outcome a coin with no financial value (the high-probability “look” stimuli) relative to the stimuli that yielded no outcome at all (Fig. S1b in the Supplementary Material). This tentative finding seems consistent with the idea that acute stress might bias behavioral responding for neutral stimuli due to augmented aberrant spontaneous phasic-dopamine release.

### 3.3. Computational modeling

To further probe the nature of the effects of acute stress on reward and punishment learning, we fitted a biologically inspired reinforcement-learning model (Frank et al., 2007) to participants’ trial-by-trial choices (see subsection 2.6.1. in “Materials and Methods” for a full description of the model). The fitted model included separate learning rates for positive (*α^+^*) and negative (*α^−^*) prediction errors, to account for the differential firing of dopaminergic neurons for positive and negative prediction errors (Schultz et al., 1997). The model also included the inverse temperature parameter, *β*, which controls the exploration/exploitation trade-off (Daw, 2011; Sutton & Barto, 1998). We then analysed the best-fitting model parameters using ANOVAs.

These analyses revealed a main effect of condition on learning rates, *F*(1, 61) = 4.69, *p* = .034, *η^2^* = 0.071, but a non-significant condition × valence interaction, *F*(1, 61) = 2.13, *p* = .15, *η^2^* = 0.034 (Fig. 3a). For completeness we performed post-hoc paired *t*-tests. Post-hoc tests revealed that *α^+^* was significantly lower in the stress condition (*M* = 0.40, *SEM* = 0.033) than in the control condition (*M* = 0.51, *SEM* = 0.038), *t*(61) = −2.25, *p* = .028, *d* = −0.29, 95% CI = [−0.21, −0.013], while *α^−^* was not significantly different between the stress (*M* = 0.25, *SEM* = 0.025) and control conditions (*M* =0.27, *SEM* = 0.027), *t*(61) = −0.72, *p* = .47, *d* = −0.092, 95% CI = [−0.098, 0.046] (Fig. 3a). Additionally, as for task-performance analyses, we repeated all statistical analyses excluding the three participants that performed below chance levels. Exclusion of these three participants revealed that the condition × valence interaction reached significance, *F*(1, 58) = 4.61, *p* = 0.036, *η^2^* = 0.074), such that *α^+^* was significantly lower in the stress condition than in the control condition, *t*(58) = −2.48, *p* = .016, *d* = −0.32, 95% CI = [−0.22, −0.024], and *α^−^* did not differ significantly between conditions *t*(58) = −0.16, *p* = .88, *d* = −0.021, 95% CI = [−0.073, 0.063]. Thus, our findings suggest that acute stress selectively decreases *α^+^*.

**Fig. 3.**
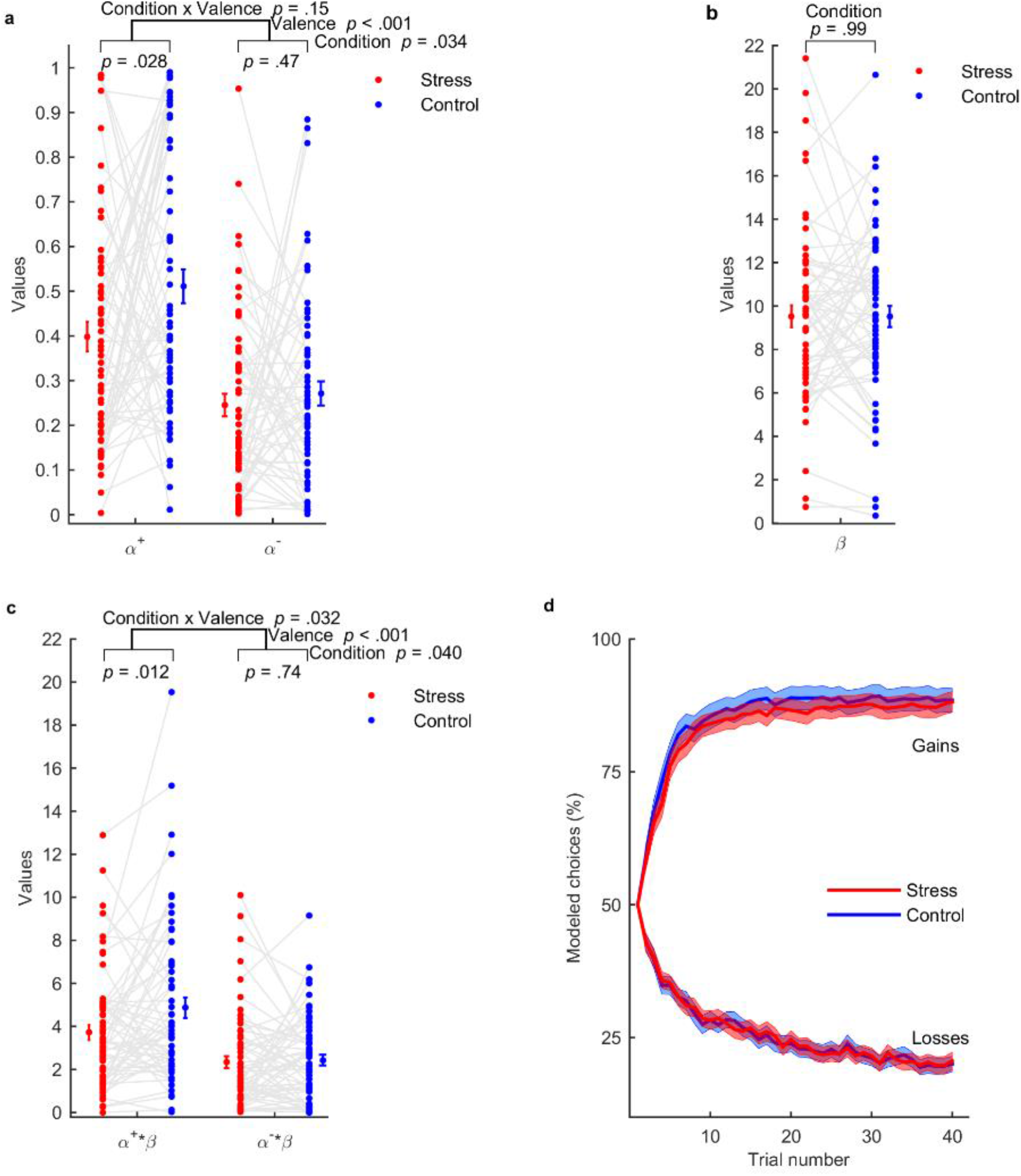
Model-fitting results. **(a)** Participants (n = 62) had lower learning rates (*α^+^* and *α^−^*) in the stress condition (red) compared with the control condition (blue). Post-hoc tests revealed a significantly lower learning rate for positive prediction errors (*α^+^*) in the stress condition comparatively to the control condition. **(b)** The inverse temperature (*β*) did not differ between conditions. **(c)** *α^+^*β*, but not *α^−^*β*, was significantly reduced in the stress condition compared with the control condition. In panels a, b and c, connected dots represent data points from the same participant. The error bar displayed on the side of the scatter plots indicate the sample mean ± standard error of the mean. **(d)** The probabilities of choosing the “correct” gain (upper part of the graph) and the “incorrect” loss (lower part of the graph) stimuli estimated by the reinforcement-learning model followed the same pattern of the actual observed choices (compare the overlap of the curves related to the stress and control conditions depicted here with the overlap of the curves that represent the actual observed choices depicted in Fig. 2c). The central lines represent the means and the filled areas represent the ± standard errors of the means.

The parameter *β*, which controls the amount of exploration/exploitation in choice selection, also did not differ significantly between the stress (*M* = 9.52, *SEM* = 0.49) and control (*M* = 9.53, *SEM* = 0.51) conditions, *t*(61) = −0.016, *p* = .99, *d* = 0.0020, 95% CI = [−0.98, 0.99] (Fig. 3b).

We further tested whether stress had a differential effect on reward compared with punishment learning by analysing the products between each learning rate (*α^±^*) and the inverse temperature (*β*). In reinforcement-learning models, *α^±^* and *β* tend to be inversely coupled (Daw, 2011; Supplementary Material of Maia & Conceição, 2017) because *α^±^* multiply by state-action values and the state-action values themselves are multiplied by *β* to compute choice probabilities (see equations in subsection 2.6.1). As a result, the parameters viewed separately can have larger estimation errors, while their product tends to be more reliably estimated (Daw, 2011; Schonberg et al., 2007), and thus better recovered. Note that the products *α^±^*β* control how strongly the outcomes impact subsequent choice preferences (Schonberg et al., 2007). Statistical analyses revealed a significant condition × valence interaction, *F*(1, 61) = 4.85, *p* = 0.032, *η^2^* = 0.074, meaning that acute stress significantly decreased *α^+^*β* (stress: *M* = 3.72, *SEM* = 0.34, control: *M* = 4.87, *SEM* = 0.48), *t*(61) = −2.58, *p* = .012, *d* = −0.33, 95% CI = [−2.04, −0.26], while not significantly affecting *α^−^*β* (stress: *M* = 2.33, *SEM* = 0.28, control: *M* = 2.44, *SEM* = 0.25), *t*(61) = −0.33, *p* = .74, *d* = −0.042, 95% CI = [−0.71, 0.51] (Fig. 3c). Exclusion of the three participants that performed below chance levels did not change the significance of the results for the parameter β, *t*(58) = 0.15, *p* = .88, *d* = 0.020, 95% CI = [−0.94, 1.10], nor for *α±*β* condition × valence interaction, *F(1*, 58) = 4.45, *p* = 0.039, *η^2^* = 0.071. Notably, the previous results were fully replicated when comparing the estimated parameters between the stress and control conditions using non-parametric Wilcoxon signed-rank, rather than parametric, post-hoc tests (*α^+^*: Z = −2.22, *p* = .026; *α^−^*: Z = −0.17, *p* = .86; *β*: Z = −0.11, *p* = .91; *α^+^*β:* Z = −2.19, *p* = .028; *α^−^*β*: Z = −0.35, *p* = .73).

To validate the used reinforcement-learning model, we confirmed that the probability of choices estimated under the reinforcement-learning model had a close correspondence with the actual observed choices across trials in both conditions, Pearson’s *r* > 0.76, *p* < .001, Spearman’s *r* > 0.70, *p* < .001 (Fig. 3d). Next, to further validate that our previous model-fitting results were reliable, we conducted parameter-recovery analyses. Those parameter-recovery analyses demonstrated that the results of our model-fitting procedure were robust both in the stress and control conditions, Pearson’s *r* > 0.67, *p* < .001, Spearman’s *r* > 0.69, *p* < .001 (see Fig. S2 in the Supplementary Material). Additionally, Bayesian model averaging analyses (Hoeting, Madigan, Raftery, & Volinsky, 1999) using both the aforementioned, neurobiologically inspired reinforcement-learning model (with separate learning rates for positive and negative prediction errors) and a nested, alternative candidate model (with a single learning rate) provided further support for the robustness of the parameters estimated by the model with separate learning rates (see Fig. S3 and “Model comparison and Bayesian model averaging” in the “Supplementary Analyses” section of the Supplementary Material).

Taken together, these computational findings indicate that under acute stress participants incorporated positive prediction errors at a lower rate, which seems to explain why acute stress impaired behavioral performance towards monetary gains during the reinforcement-learning task.

## 4. Discussion

Acute stress is present in day-to-day life, and people recurrently need to make choices and learn from the rewarding or punishing outcomes of those choices whilst under stress. In this study we investigated whether and how acute stress impacted reward and punishment learning in men using a reinforcement-learning framework. We hypothetized that acute stress would impair reward, and possibly punishment, learning by reducing learning rates for positive and negative prediction errors, respectively. We found that acute stress impaired behavioral performance towards monetary gains, but not losses, and that this impaired performance was explained by a decreased learning rate for positive prediction errors.

### 4.1. Effect of acute stress on reward learning

Our finding that acute stress impaired reward-seeking performance is consistent with previous several studies which found reduced reward responsiveness under acute stress (Bogdan et al., 2010, 2011; Bogdan & Pizzagalli, 2006; Morris & Rottenberg, 2015; Paret & Bublatzky, 2020), particularly in high stress-reactive individuals (Berghorst et al., 2013). At a first sight, however, some of the extant literature may seem equivocal, possibly due to critical methodological differences related to stress operationalization (Porcelli & Delgado, 2017). A significant number of studies have investigated the effects of stress on learning using different paradigms, such as the cold pressor test (Byrne et al., 2019; Ehlers & Todd, 2017; Glienke, Wolf, & Bellebaum, 2015; Lighthall et al., 2013; Otto et al., 2013; Paul, Bellebaum, Ghio, Suchan, & Wolf, 2019) or the Trier social stress test (Boyle, Stanton, Eisenberger, Seeman, & Bower, 2019; Kruse et al., 2018; Petzold et al., 2010; Radenbach et al., 2015), in which acute stress is induced before the learning task. In these paradigms, stress induction precedes any learning processes, thus the stress-induced emotional state may be less concurrent with the cognitive processes that operate during the task. As such, it is unclear whether these paradigms probe the effects of acute stress or of recovery from stress on learning (Hermans, Henckens, Joëls, & Fernández, 2014). Critically, studies that induce stress before the learning task may suggest divergent behavioral results from ours (e.g., Byrne et al., 2019; Lighthall et al., 2013), whereas studies that have induced stress during the learning task (as ours) seem to be in agreement with our findings of impaired reward learning during acute-stress exposure (Berghorst et al., 2013; Bogdan et al., 2010, 2011; Bogdan & Pizzagalli, 2006; Morris & Rottenberg, 2015; Paret & Bublatzky, 2020). Therefore, cross-study comparisons suggest that individuals may perform more poorly when learning to maximize rewards whilst exposed to an acute stressor.

In this study we further inspected the computational mechanisms behind impaired reward learning under acute stress. Using a reinforcement-learning model with separate learning rates for positive and negative prediction errors (Frank et al., 2007), we found that acute stress reduced the rate at which participants learned from positive prediction errors. A reduced learning rate for positive predictions errors means that under acute stress individuals learned more slowly about unexpected rewards and therefore took longer to adapt their behavior on the basis of the more recent rewarding outcomes of their choices. Other studies have successfully applied this reinforcement-learning model to analyze data from distinct tasks to describe the genetic (Doll et al., 2011; Frank et al., 2007) and neural (Diederen et al., 2016; Lefebvre, Lebreton, Meyniel, Bourgeois-Gironde, & Palminteri, 2017; Niv, Edlund, Dayan, & Doherty, 2012) correlates of cognition and behavior. However, to the best of our knowledge, no studies had attempted to model how acute stress affects cognition, particularly reward and punishment learning, using such model. Another reinforcement-learning model commonly used in the stress literature includes distinct learning rates for gain and loss trials regardless of the valence of the prediction error (Aylward et al., 2019; Robinson et al., 2013; Treadway et al., 2017), meaning that learning from rewards in gain trials (or punishments in loss trials) is modeled the same way as learning from omission of rewards in gain trials (or omission of punishments in loss trials, respectively), which may not be so well supported by neurobiological evidence as a model that assumes distinct learning rates based on the prediction errors’ valence (as the one we used) (Maia & Frank, 2011; O’Doherty, Dayan, Friston, Critchley, & Dolan, 2003; Schultz et al., 1997). Thus, our findings seem to contribute to a better mechanistic understanding of how acute stress may impact reward learning.

Given that the (quantifiable) parameters from the reinforcement-learning model that we used seem to reflect specific dopaminergic-related neural mechanisms (e.g., Frank et al., 2007; Frank & O’Reilly, 2006), our computational findings may shed light on putative neural mechanisms underlying the impact of acute stress on cognition and behavior. Specifically, our computational findings seem broadly consistent with the proposed neurobiological account of dopaminergic neurons functioning under acute stress. Acute stress is thought to induce aberrant spontaneous dopamine release (Anstrom et al., 2009; Anstrom & Woodward, 2005; Cabib & Puglisi-Allegra, 2012; Valenti et al., 2011); stress-induced spontaneous dopamine release, in turn, may disrupt the adaptive striatal phasic-burst dopamine responses that signal positive prediction errors (Bilder et al., 2004; Daberkow et al., 2013; Grace, 2016; Maia & Frank, 2017; Werlen et al., 2020), which would explain the stress-induced impairment of reward learning. While our computational results suggest that acute stress disrupts striatal responses to prediction errors during reward learning (see also Huys et al., 2013), we cannot disregard, however, that acute stress may also affect reward learning via prefrontal cortex disturbances (Lighthall et al., 2013; Otto et al., 2013; Whitmer, Frank, & Gotlib, 2012).

### 4.2. Effect of acute stress on punishment learning

Acute stress impaired reward learning, but we found no evidence for an effect of acute stress on punishment learning. In line with this behavioral data, computational-modeling analyses also provided no evidence that acute stress affected the learning rate for negative prediction errors.

According to a long standing influential loss aversion framework (Kahneman & Tversky, 1979), losses can have more debilitative potential than gains; therefore, as an adaptive strategy, it is possible that individuals may be more attuned to losses than to gains (Lejarraga, Hertwig, & Gonzalez, 2012; Yechiam, 2019; Yechiam & Hochman, 2013), explaining why punishment learning was spared under acute stress. However, recent evidence suggests that individuals learn gain associations better than loss associations in reinforcement-learning tasks despite symmetrical task structure and symmetrical outcome probabilities (Lin, Cabrera-Haro, & Reuter-Lorenz, 2020) (as in our task). Indeed, our behavioral data indicated that participants performed better when learning to seek gains than when learning to avoid losses in both the stress and control conditions. In addition, our computational data indicated that the learning rate for positive prediction errors was significantly higher than the learning rate for negative prediction (see also Lefebvre et al., 2017) in both conditions. Thus, it seems unlikely that acute stress preserved the mechanisms involved in punishment learning, but not in reward learning, due to heightened attention towards losses relatively to gains (i.e., by preferentially decreasing attention towards gains).

One potential neurocognitive explanation for such lack of effect of acute stress on loss avoidance is that D2 dopamine receptors, which mediate punishment learning (Frank & O’Reilly, 2006; Maia & Frank, 2011), are already mostly activated at baseline dopamine levels, so their activation might be affected by decreases, but less so by increases, in dopamine levels (Maia & Conceição, 2017; Möller & Bogacz, 2019). Finally, non-dopaminergic mechanisms may also be involved in punishment learning (Boureau & Dayan, 2011; Moran et al., 2018), which may partially explain why previous studies using the same reinforcement-learning task also did not find significant effects of pharmacological manipulations of the dopaminergic system on punishment learning (Eisenegger et al., 2014; Pessiglione et al., 2006).

### 4.3. Acute stress and behavioral performance in neutral trials

We found preliminary evidence that acute stress biased behavioral responding to neutral stimuli, as the choice of the high-probability “look” stimuli — compared to the stimuli with a high probability of yielding “nothing” — was augmented under stress compared to the control condition (for a detailed discussion about this finding, see the Supplementary Material). Although very tentative, this finding might be of relevance as excessive, aberrant spontaneous dopamine release is thought to underlie increased behavioral responding and aberrant learning for neutral stimuli (Maia & Frank, 2017; Roiser et al., 2013). Still, whether acute stress promotes aberrant valuation of neutral stimuli via dopaminergic disturbances remains unknown and should be further investigated.

### 4.4. Limitations

In this study, we induced acute stress in participants, using a repetitive and uncontrollable sound, whilst they completed a reinforcement-learning task. We exposed participants to the sound during the learning task to ensure that acute stress was contingent on the learning processes, but at the expense of possibly confounding the induction of stress with distraction. To avoid this potential confound, we used a sound that was always constant and repetitive, as evidence suggests that rare and unexpected changes in an otherwise repetitive sound sequence seem to induce distraction more robustly (Hughes, 2014; Parmentier, 2014; Parmentier, Elford, Escera, Andrés, & Miguel, 2008). Furthermore, if the sound was acting as a distractor, rather than as a stressor, we would expect a general behavioral impairment rather than a selective effect in gains. Relatedly, if the manipulation had acted mostly as a distractor, we would expect participants to behave more at random (Tsushima & Nakayama, 2010), which would likely be captured in a reduction of the inverse temperature parameter — reflecting increased random behavior — in the stress condition.

Finally, only men were included in this study, to avoid the potential confounding effects of menstrual-cycle-dependent variation on stress responsivity (Ossewaarde et al., 2010) and reward and punishment processing (Diekhof et al., 2020; Dreher et al., 2007). Our finding that acute stress disrupts reward learning in men seems to be in line with previous reports showing that acute stress disrupts reward-seeking behavior in women (Berghorst et al., 2013; Bogdan et al., 2010, 2011; Bogdan & Pizzagalli, 2006; Morris & Rottenberg, 2015; Paret & Bublatzky, 2020), but further studies are needed to assess whether acute stress has the same computational effects on reward and punishment learning in men and women.

## 5. Conclusion

We present evidence that acute stress reduces how quickly male adults integrate the unexpected rewarding outcomes of their choices over time. Our results are consistent with a neurobiological framework of stress-induced dopaminergic disturbances and can thus contribute to a better understanding of the computational mechanisms that underlie the deleterious impact of acute stress on reward learning. Ultimately, this study might offer key mechanistic insights into the impact of acute stress in everyday life.

## Author contributions

J. Carvalheiro, A. Mesquita and A. Seara-Cardoso developed the study concept with critical input from V. Conceição. Testing and data collection were performed by J. Carvalheiro under the supervision of A. Mesquita and A. Seara-Cardoso. Data analyses and interpretation was carried out by J. Carvalheiro and V. Conceição. J. Carvalheiro drafted the manuscript. All authors revised and edited the manuscript.

## Acknowledgments

We thank Maël Lebreton for helpful discussions. This work was supported by grants from the Portuguese Foundation for Science and Technology (FCT) to A. Seara-Cardoso [PTDC/MHC-PCN/2296/2014, co-financed by FEDER through COMPETE2020 under the PT2020 Partnership Agreement (POCI-01-0145-FEDER-016747)] and to A. Mesquita (IF/00750/2015). J. Carvalheiro was supported by a FCT PhD fellowship (PD/BD/128467/2017). This study was conducted at the Psychology Research Centre (PSI/01662), School of Psychology, University of Minho, supported by FCT and the Portuguese Ministry of Science, Technology and Higher Education (UID/PSI/01662/2019), through national funds (PIDDAC).

