## Supplementary material for "Acute stress impairs reward learning in men": SI

This Supplementary Material includes:

Supplementary Analyses

Supplementary Figures (Fig. S1 to S3)

Supplementary Tables (Tables S1 to S4)

Supplementary References

### Supplementary Analyses

#### Analysis of neutral trials

Evidence from non-human animal studies suggests that acute stress increases aberrant, spontaneous phasic-dopamine release (Anstrom, Miczek, & Budygin, 2009; Anstrom & Woodward, 2005; Cabib & Puglisi-Allegra, 2012; Valenti, Lodge, & Grace, 2011). In turn, excessive spontaneous phasic-dopamine release is thought to drive increased behavioral responding and aberrant learning for neutral stimuli (Boehme et al., 2015; Maia & Frank, 2017; McCutcheon, Abi-Dargham, & Howes, 2019). With this in mind, we conducted exploratory analyses to test whether acute stress biased participants' responses during the neutral trials of the reinforcement-learning task (Murray et al., 2008). Each neutral trial consisted of a choice between two different stimuli, which were either associated with a high (0.8), or with a low (0.2), probability of observing as outcome: 1) "look" and a 0.5€ coin (but the participants did not win nor lose money), or 2) "nothing" and a gray square (participants also did not win nor lose money; Fig. S1a). Note that the "look" outcome was only present in neutral trials, whereas the "nothing" outcome was also present in gain and loss trials (see Fig. 1 in the main manuscript for a depiction of gain and loss trials). Although none of the neutral stimuli led to reinforcement (i.e., none of the stimuli was associated with financial outcomes), one of the stimuli was associated, at a high probability, with the observation of a coin ("look"). Aberrant, spontaneous phasic release of dopamine during coin observation ("look") could conceivably act similarly to the phasic release of dopamine that signals prediction errors (Hyland, Reynolds, Hay, Perk, & Miller, 2002) in gain trials following the observation of the 0.5€ coin. Thus, we asked whether stress-induced spontaneous release of dopamine would lead to a behavioral bias during neutral trials towards the selection of the stimulus that led more often to the coin ("look").

To explore the impact of acute stress on behavioral responding during neutral trials, we conducted a generalized linear-mixed effects model with a binomial distribution of the response variable (0 or 1, for the selection of the stimuli with low and high probabilities of obtaining the "look" outcome, respectively; "logit" link function). Predictor variables included condition (stress or control), block (1 or 2), and trial number (from 1 to 40). The model was fitted using Matlab's *fitglme* function. Importantly, we observed a significant main effect of condition on the selection of the high-probability "look" stimuli, $\beta = 0.18$, $p < .001$, 95% confidence interval = [0.096, 0.26] (Fig. S1b and Table S2). Specifically, participants exhibited a bias towards selecting the high-probability "look" stimuli during the stress condition (% choices: $M = 50.0$, $SEM = 2.8$) compared with the control condition (% choices: $M = 46.0$, $SEM = 2.7$). This result suggests that acute stress modulates behavioral responding for neutral stimuli, possibly due to stress-induced spontaneous phasic-dopamine release. We cannot discard, however, other alternative interpretations for this tentative finding.

**Model comparison and Bayesian model averaging**

To confirm that the reinforcement-learning model that we used (Frank, Moustafa, Haughey, Curran, & Hutchison, 2007) provided a robust account of the data, we also tested a nested variant of that model. Non-human animal and human studies suggest that dopaminergic neurons are differentially implicated in the coding of positive and negative prediction errors (Frank et al., 2007; Hikida, Kimura, Wada, Funabiki, & Nakanishi, 2010; Schultz, Dayan, & Montague, 1997) and that learning from positive and negative prediction errors likely occurs via the differential effects of dopamine on the plasticity of corticostriatal synapses (Collins & Frank, 2013; Maia & Conceição, 2017; Möller & Bogacz, 2019); such neurobiological evidence motivated us to use a model with separate learning rates for positive ($\alpha^+$) and negative ($\alpha^-$) prediction errors (double-$\alpha$ model) [for examples of studies using this model, see Diederen et al., (2016); Frank et al., (2007); Horga et al., (2015); Lefebvre, Lebreton, Meyniel, Bourgeois-Gironde, & Palminteri, (2017); Niv, Edlund, Dayan, & Doherty, (2012)]. In the classical reinforcement-learning framework, however, such distinction between positive and learning rates is not common (Sutton & Barto, 1998). Therefore, the second model that we used was the standard Q-learning model with a single learning rate ($\alpha$) for both positive and negative prediction errors (single-$\alpha$ model) (Sutton & Barto, 1998). Both models included the inverse temperature parameter ($\beta$), which controls for the amount of exploration/exploitation in choice selection (Daw, 2011; Sutton & Barto, 1998). Given that these two models could show a divergent quality of fits for the stress and control conditions, we fitted both models (double-$\alpha$ and single-$\alpha$) to the stress and control conditions independently. Then, we compared the two models using random-effects Bayesian model comparison (Rigoux, Stephan, Friston, & Daunizeau, 2014; Stephan et al., 2009).

Bayesian model comparison primarily relies on the model evidence, which quantifies how likely the observed data is under a given model (Daw, 2011). For standard reinforcement-learning models, however, the calculation of the corresponding model evidences is not tractable (Daw, 2011), so approximations must be used. Two frequently used approximations are the Bayesian information criterion (BIC) and the Akaike's information criterion (AIC). The BIC and AIC are relatively poor approximations of the model evidence and their use commonly leads to the selection of different models, given that the BIC and AIC typically favor simpler and more complex models, respectively (Penny, Stephan, Mechelli, & Friston, 2004), especially when there is a considerable number of observations (or trials), $n$:

$$\mathrm{BIC} = \log\big(P(D|M, \theta_M)\big) - \frac{p}{2}\log(n),$$

$$\text{AIC} = \log\left(P(D|M, \theta_M)\right) - p,$$

where $P(D|M, \theta_M)$ is the likelihood of the data $D$ (in our case, the observed responses from each subject in each condition) given the model $M$ and parameter values $\theta_M$, and $p$ is the number of free parameters of the model. Note that, in the equations above, the BIC and AIC are approximations to the log model evidences, not to the model evidences.

We compared the two models between conditions and at the single condition level, using random-effects Bayesian model comparison (Rigoux et al., 2014; Stephan et al., 2009) as implemented in the Variational Bayesian Analysis (VBA) toolbox (Daunizeau, Adam, & Rigoux, 2014). In the VBA toolbox, the input data to the Bayesian model comparison procedure was composed of the log-model-evidence approximations (BIC or AIC) obtained for each subject, in each condition, for each model (double-$\alpha$ and single-$\alpha$). This procedure allowed estimating the model frequencies, and the exceedance probabilities, of each model within the tested set of candidate models, given the data gathered from all subjects. The estimated model frequencies quantify the prevalence of a model in the population. The exceedance probability (EP) measures how likely it is that any given model is more frequent than all other models in the comparison set (Stephan et al., 2009). Here, we report protected exceedance probabilities (PEPs), which correct exceedance probabilities for the possibility that observed differences in model evidences (across groups or conditions) are due to chance (Rigoux et al., 2014). A PEP above 0.95 provides strong evidence for the selection of a given candidate model (Rigoux et al., 2014).

Bayesian model comparison revealed divergent results when using the different approximations to the log model evidence (i.e., the BIC and AIC; Table S3). Model comparison using the BIC did not lead to confident selection of the same model across the stress and control conditions (PEP = 0.37), nor to confident selection of different models in each condition (stress condition: PEP double-$\alpha$ = 0.04, PEP single-$\alpha$ = 0.96; control condition: PEP double-$\alpha$ = 0.29, PEP single-$\alpha$ = 0.71). Similarly, model comparison using the AIC was inconclusive, as it also did not lead to selection of the same model across the stress and control conditions (PEP = 0.92), nor to the selection of different models in each condition (stress condition: PEP double-$\alpha$ = 0.57; PEP single-$\alpha$ = 0.43; control condition: PEP double-$\alpha$ = 0.995, PEP single-$\alpha$ = 0.005). Thus, we turned to Bayesian model averaging (BMA) (Fragoso, Bertoli, & Louzada, 2018; Hoeting, Madigan, Raftery, & Volinsky, 1999), which can be used to combine parameter estimates from multiple nested models, when no single model clearly is best. Specifically, for each approximation to the log model evidence, we calculated weighted point parameter estimates for each subject and condition, by multiplying the subject-condition-specific parameter estimates obtained through each reinforcement-learning model (double-$\alpha$ and single-$\alpha$) by the posterior model probabilities obtained with each approximation (BIC and AIC) for each subject in each condition.

To confirm that the effects of acute stress on the estimated parameters could be successfully recovered independently of the used approximations to the log model evidence, we analyzed the parameters estimated through BMA when using either the BIC or the AIC. The ensuing analyses demonstrated that, regardless of the used approximation (BIC and AIC; Fig. S3), we were able to recapitulate the effects of acute stress on the estimated parameters reported in the main text, when using only the model with separate learning rates for positive and negative prediction errors (compare Fig. S3 with Fig. 3a-c in the manuscript), except for the interaction condition × valence in $\alpha^{\pm}*\beta$ when the BIC was used as an approximation to log model evidence (Fig. S3e). Still, when we applied post-hoc tests to the BMA results, we were always able to replicate the differences observed in the original parameters between conditions (again, compare

Fig. S3 with Fig. 3a-c in the manuscript). Additionally, all parameters estimated through BMA were highly correlated with those used in the main paper analyses (Pearson's $r \geq 0.62$, $p < .001$, Spearman's $r > 0.62$, $p < .001$). In sum, although model comparison using the BIC or AIC was not conclusive *per se*, BMA results were always in strikingly agreement with those obtained when using only the model with separate learning rates for positive and negative prediction errors.

### Supplementary Figures

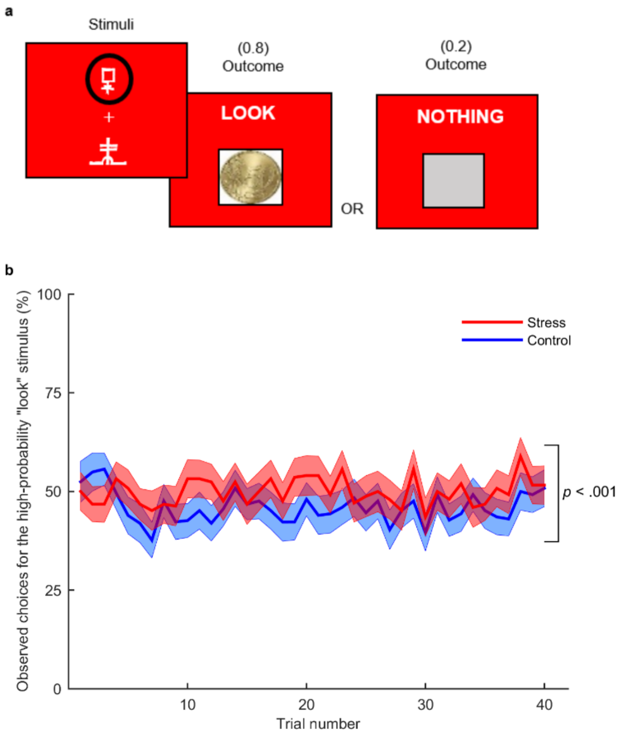


**Fig. S1.** Neutral trials and behavioral performance in neutral trials. **(a)** Each neutral stimulus was associated with a certain probability (0.8 or 0.2) of obtaining the outcome “look” and a 0.5€ coin—the obtained monetary outcome was zero, meaning that participants did not win or lose money—and with the reciprocal probability (0.2 or 0.8) of obtaining “nothing” and a gray square. Here, we depict an example for the stress condition in which the participant selected the stimulus with 0.8 probability of obtaining the outcome “look” (i.e., the high-probability “look” stimulus). **(b)** Participants chose the stimulus associated at a higher probability with the “look” at the coin outcome significantly more often in the stress condition (red) than in the control condition (blue). The curves represent the trial-by-trial percentage of participants (n = 62) who chose the high-probability “look” stimulus in the stress and control conditions, averaged across the two blocks in each condition. The central lines represent the means and the filled areas represent ± standard errors of the mean.

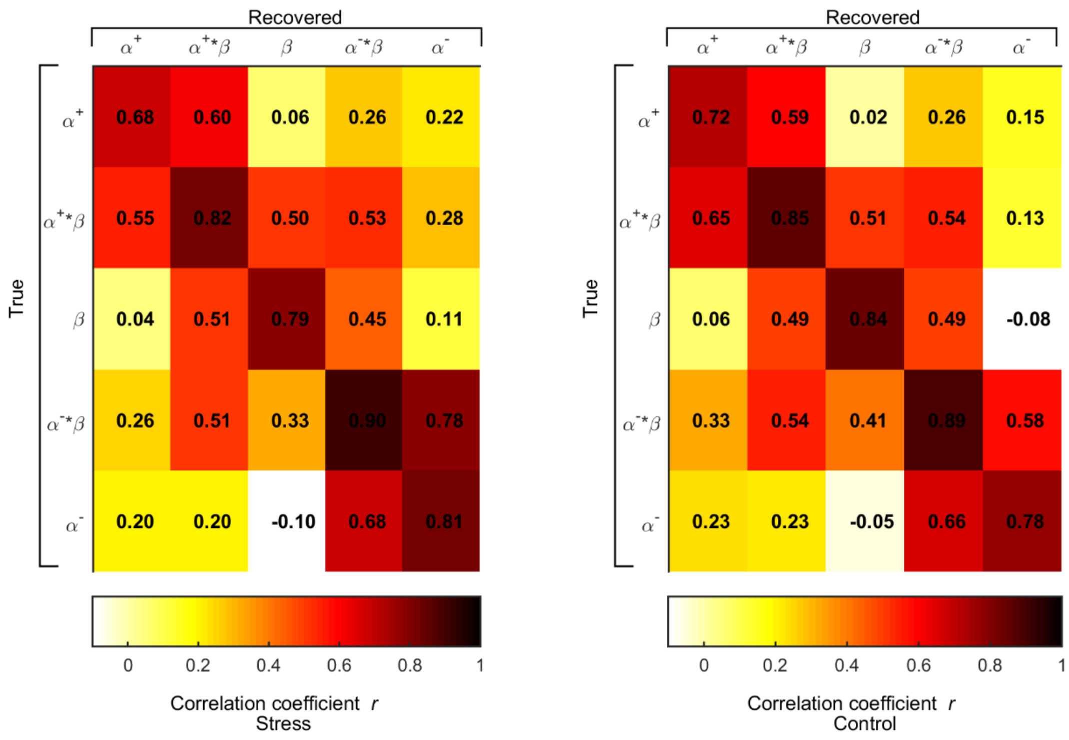


**Fig. S2.** Correlation matrices between the estimated and recovered parameters averaged across 100 simulations in the stress (left) and control (right) conditions. The matrices include the values of Pearson's *r* for the correlations between the parameters that were used to generate the simulated data ("True") and those that were obtained by applying the parameter-estimation procedure to the simulated data ("Recovered"). Dark red values of *r* indicate a strong correlation; dark-red diagonal values indicate therefore that parameter recovery was "successful".

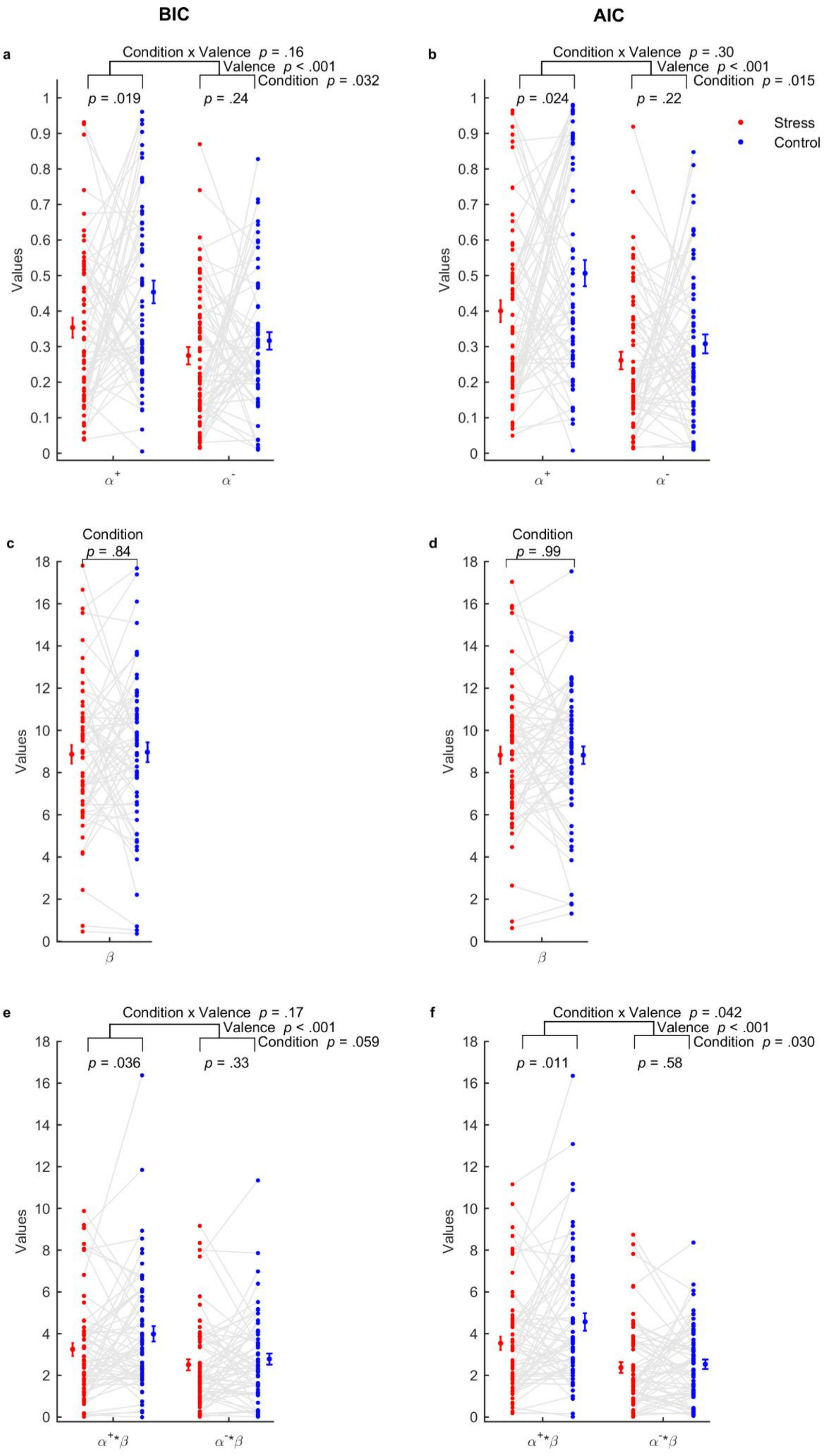
BIC
AIC
a
Condition x Valence p = .16
Valence p < .001
Condition p = .032
p = .019
p = .24
b
Condition x Valence p = .30
Valence p < .001
Condition p = .015
p = .024
p = .22
Stress
Control
Values
α+
α-
c
Condition
p = .84
d
Condition
p = .99
β
e
Condition x Valence p = .17
Valence p < .001
Condition p = .059
p = .036
p = .33
f
Condition x Valence p = .042
Valence p < .001
Condition p = .030
p = .011
p = .58
α+*β
α-*β

**Fig. S3.** Parameters estimated using Bayesian model averaging (BMA) across the double-$\alpha$ and single-$\alpha$ models, when using either the Bayesian information criterion (BIC; left panels) or the Akaike information criterion (AIC; right panels) as approximations to the log model evidence, in the stress (red) and control (blue) conditions. **(a, b)** Learning rates for positive, $\alpha^{+}$, and negative, $\alpha^{-}$, prediction errors. **(c, d)** Inverse temperature parameter, $\beta$. **(e, f)** Products between each learning rate and $\beta$, $\alpha^{+} * \beta$ and $\alpha^{-} * \beta$. Connected dots represent individual data points in the within-subject design (n = 62). The error bars displayed on the sides of the scatter plots indicate the sample means ± standard errors of the means.

#### Supplementary Tables

**Table S1**. Analyses of behavioral performance during the reinforcement-learning task. Estimated fixed effects coefficients (± standard errors) from generalized linear mixed-effects models. The results reported in the main text refer to a model that included all main effects and the interaction of interest (condition × valence). The significance of the condition × valence interaction did not change when including the other two-factor interactions in the model. As an additional check, we included the significant two-factor interactions ($p < .05$) in a three-factor interaction model. Subject identity was modeled via random intercepts. The reference levels were the control condition, first block, and loss valence.

| Fixed effects | Coefficient | Standard error | *T* | *p* | 95% Confidence interval |
|---|---|---|---|---|---|
| Intercept | 0.32 | 0.11 | 2.96 | .003 | [0.11, 0.58] |
| Trial | 0.048 | 0.0025 | 18.9 | < .001 | [0.043, 0.053] |
| Block | 0.65 | 0.057 | 11.4 | < .001 | [0.54, 0.76] |
| Condition | 0.048 | 0.089 | 0.54 | .59 | [-0.12, 0.22] |
| Valence | 0.33 | 0.056 | 5.94 | < .001 | [0.22, 0.45] |
| Condition × Trial | 0.0027 | 0.0035 | 0.77 | .44 | [-0.0042, 0.0097] |
| Condition × Block | -0.26 | 0.078 | 2.96 | < .001 | [-0.42, -0.11] |
| Condition × Valence | -0.19 | 0.078 | -2.37 | .018 | [-0.34, -0.032] |
| Condition × Valence × Block | -0.088 | 0.16 | -0.55 | .58 | [-0.40, 0.22] |

| | | | | | |
|---|---|---|---|---|---|
| Condition × Trial × Block | -0.014 | 0.0072 | -1.90 | .057 | [-0.028, 0.00041] |
| Condition × Trial × Valence | -0.0042 | 0.0072 | -0.59 | .55 | [-0.018, 0.0098] |

**Table S2**. Analyses of behavioral performance during neutral trials. Estimated fixed-effects coefficients (± standard errors) from generalized linear mixed-effects models. The final model included all main effects. Subject identity was modeled via random intercepts. The reference levels were the control condition, first block, and loss valence.

| Fixed effects | Coefficient | Standard error | *t* | *p* | 95% Confidence interval |
|---|---|---|---|---|---|
| Intercept | 0.07 | 0.11 | 0.65 | .51 | [-0.15, 0.30] |
| Trial | 0.000055 | 0.0018 | 0.030 | .97 | [-0.0036, 0.0037] |
| Block | -0.48 | 0.043 | -11.33 | < .001 | [-0.57, -0.40] |
| Condition | 0.18 | 0.043 | 4.21 | < .001 | [0.096, 0.26] |
| Condition × Trial | 0.0034 | 0.0037 | 0.93 | .35 | [-0.0038, 0.011] |
| Condition × Block | 0.11 | 0.085 | 1.25 | .21 | [-0.061, 0.27] |

**Table S3.** Results from Bayesian model comparison between the reinforcement-learning model used in the main text (double-α) and its nested model with a single learning rate (single- α). NP: number of parameters; M: mean; SEM: standard error of the mean; BIC: Bayesian information criterion; AIC: Akaike information criterion; EF: estimated model frequencies; PEP: protected exceedance probability.

| | | | BIC | | | AIC | | |
|---|---|---|---|---|---|---|---|---|
| Condition | Model | NP | *M* ± *SEM* | EF | PEP | *M* ± *SEM* | EF | PEP |
| Stress | Double-α | 3 | -67.6 ± 2.8 | 0.27 | 0.04 | -63.0 ± 2.8 | 0.61 | 0.57 |
| | Single-α | 2 | -67.5 ± 2.9 | 0.73 | 0.96 | -64.4 ± 2.9 | 0.39 | 0.43 |
| Control | Double-α | 3 | -63.8 ± 2.9 | 0.34 | 0.29 | -59.2 ± 2.9 | 0.96 | 0.995 |
| | Single-α | 2 | -64.2 ± 3.1 | 0.66 | 0.71 | -61.2 ± 3.1 | 0.04 | 0.005 |
